# Plasticity induction in the ventromedial prefrontal cortex during REM sleep improves fear extinction memory consolidation

**DOI:** 10.1101/2025.03.31.646182

**Authors:** Vuk Marković, Gaetano Rizzo, Fatemeh Yavari, Carmelo M. Vicario, Michael A. Nitsche

## Abstract

Anxiety disorders (ADs) are among the most prevalent mental health conditions, yet first-line treatments often yield only moderate effectiveness. The fear conditioning paradigm is commonly used to investigate fear and extinction learning, revealing deficits in these processes and dysfunctional activity in the ventromedial prefrontal cortex (vmPFC) and amygdala in individuals with ADs. The vmPFC plays a critical role in regulating activity of the amygdala and consolidation of fear extinction memory. Notably, transcranial direct current stimulation (tDCS) has shown promise in enhancing fear extinction by modulating vmPFC activity. Additionally, rapid eye movement (REM) sleep has been suggested to be crucial for fear extinction memory consolidation.

This study investigated the role the vmPFC during REM sleep in fear extinction memory consolidation. Thirty-two participants underwent a 3-day differential fear conditioning paradigm, with tDCS or sham stimulation applied during REM sleep over the vmPFC. Outcome measures included skin conductance responses (SCR) and subjective ratings of arousal, fear, and valence. Results indicate that tDCS during REM sleep enhances fear extinction memory consolidation, as measured by SCR. Furthermore, participants reported an increased subjective arousal following tDCS. These findings suggest that tDCS during REM sleep may hold potential for improving exposure-based treatments for ADs by strengthening fear extinction memory.

## 1. Introduction

Anxiety disorders (ADs) are among the most prevalent mental health conditions, affecting over 300 million people globally. ADs impose a significant impact on individual well-being and lead to substantial economic costs (Gustavsson et al., 2011; GBD 2019 Mental Disorders Collaborators, 2022; Xiong et al., 2022). Despite the widespread use of first-line treatments such as selective serotonin reuptake inhibitors (SSRIs) and cognitive-behavioral therapy (CBT), these approaches offer only moderate success, with 40% of individuals not responding to treatment and around 20% of the patients dropping out from treatment strategies prematurely, underscoring the need for new therapeutic strategies (Bystritsky, 2006; Taylor et al., 2012; Fernandez et al., 2015; Bandelow et al., 2017; Jakubovski et al., 2019).

The fear conditioning and extinction paradigm is a well-established experimental model to understand the mechanisms underlying fear and extinction learning (Duits et al., 2015; Lonsdorf et al., 2017; Bouton et al., 2021). In a typical differential fear conditioning protocol, one stimulus (CS+, signaling danger) is paired with an aversive stimulus (US), while another stimulus (CS-, signaling safety) is not, enabling to assess responses towards threat, but also safety. During extinction, the CS+ is no longer paired with the aversive stimulus leading to inhibition of fear responses.

Individuals with ADs and related disorders, such as post-traumatic stress disorder (PTSD), often exhibit dysfunctional fear acquisition and extinction. PTSD patients show enhanced fear responses during fear acquisition to the CS+ compared to healthy controls (Norrholm et al., 2011). Moreover, individuals with ADs demonstrate increased fear responses to the CS-during fear acquisition (Duits et al., 2015; Rabinak et al., 2017), and continue to exhibit elevated fear responses to the CS+ during extinction (Duits et al., 2015).

The neural circuits underlying fear acquisition and extinction involve a broad network of brain regions, including the amygdala, insula, hippocampus, anterior cingulate cortex (ACC), and dorsolateral (dlPFC) and ventromedial prefrontal cortex (vmPFC) (for detailed reviews, refer to Sehlmeyer et al., 2009; Milad & Quirk, 2012; Fullana et al., 2016, 2018; Bouton et al., 2021). Here, the amygdala and vmPFC are crucial for processes involved in acquiring and extinguishing fear responses, including neuroplasticity.

The amygdala plays a key role in forming fear and extinction memories and eliciting fear responses. During fear acquisition, the basolateral nucleus of the amygdala (BLA) integrates sensory inputs from the CS and US, transmitting outputs to the central nucleus (CEA) to trigger fear responses (LeDoux, 2007; Izquierdo et al., 2016). Furthermore, the intercalated nucleus of the amygdala (ITC) receives input from the BLA and exerts inhibitory control over neurons in the CEA, a mechanism essential for fear extinction (Likhtik et al., 2008). Neuroimaging studies support this involvement of the amygdala in fear learning and extinction, showing increased activity during fear acquisition and decreased activity during extinction (Phelps et al., 2004; Sehlmeyer et al., 2009).

The vmPFC, which is a homolog of the infralimbic cortex (IL) in rodents, is crucial for fear extinction and retrieval. It modulates the CEA output through its influence on ITC and BLA neurons, contributing to the suppression of fear responses during extinction (Quirk & Mueller, 2008; Do-Monte et al., 2015; Kim et al., 2016). Human neuroimaging studies corroborate the role of the vmPFC in fear extinction (Phelps et al., 2004; Kalisch et al., 2006; Milad et al., 2007; Fullana et al., 2018).

Deficient amygdala and vmPFC functions have been observed in individuals with ADs and PTSD. For example, hyperactivity of the amygdala during fear acquisition (Bremner et al., 2005; Marin et al., 2017) and extinction (Milad et al., 2009), as well as hypoactivation of the vmPFC during extinction recall (Marin et al., 2017), have been reported. Additionally, disrupted functional connectivity between the amygdala and vmPFC suggests reduced emotional regulation, further implicating the relevance of these regions for the development and maintenance of ADs (Hilbert et al., 2014; Milad et al., 2012, 2014).

REM sleep plays a critical role in fear extinction memory consolidation. REM sleep deprivation in rodents impairs fear extinction and recall (Silvestri, 2005; Fu et al., 2007; Jung & Noh, 2021) and inhibiting the IL during REM sleep disrupts extinction memory consolidation (Hong et al., 2024). In humans, increased vmPFC activation and enhanced extinction recall have been observed following REM sleep (Spoormaker et al., 2010; Menz et al., 2016). Brain regions activated during REM sleep, such as the amygdala, hippocampus, ACC, and vmPFC, overlap with those involved in fear learning and extinction, highlighting the importance of REM sleep in these processes (Van Der Helm et al., 2011; Pace-Schott et al., 2015).

Transcranial direct current stimulation (tDCS) has emerged as an innovative tool for modulating brain functions. This technique involves applying a small electrical current over the scalp to modulate cortical excitability in a polarity-dependent manner—anodal stimulation enhances excitability, while cathodal stimulation reduces it with standard protocols at the macroscale level (Nitsche & Paulus, 2000; Nitsche et al., 2008). In addition, application of tDCS for a couple of minutes leads to LTP- and LTD-like plasticity, depending on the polarity of stimulation (Stag et al., 2018). When applied over the vmPFC, anodal tDCS has been shown to enhance fear extinction and recall, likely by inducing LTP-like plasticity (Vicario et al., 2019, 2020; Marković et al., 2021; Adams et al., 2023; Boehme et al., 2024).

tDCS has potential for enhancing memory consolidation during sleep. Anodal tDCS during REM sleep has been shown to enhance motor memory consolidation (Nitsche et al., 2010). Similarly, anodal slow-oscillatory tDCS during non-REM sleep enhanced declarative memory consolidation (Marshall et al., 2004, 2006; Ladenbauer et al., 2016; Cellini et al., 2019).

To the best of our knowledge, no studies have directly assessed the causal role of REM sleep in fear extinction memory consolidation via non-invasive brain stimulation. We hypothesize that inducing LTP-like plasticity with tDCS in the vmPFC during REM sleep will enhance fear extinction memory consolidation by enhanced synaptic strengthening. The results of the study are expected to provide important insights into the potential of tDCS for enhancing treatment efficacy of ADs.

## 2. Methods

### 2.1. Participants

The study employed a single-blind, randomized, sham-controlled design. All participants were blinded to the tDCS condition. A total of 50 subjects were recruited, with 18 participants dropping out or being excluded for various reasons: 8 subjects failed to achieve a sufficient REM sleep duration, 3 did not exhibit adequate fear acquisition skin conductance responses (SCR) (average of all CS+ SCR > average of all SCR CS-), 2 discontinued the experiment during the sleep phase due to discomfort, 1 subject had abnormal electrodermal activity (SCR > 3 standard deviations (SD) from the mean), 1 due to equipment failure, 1 did not attend the third day of the study, 1 did not have the mandatory complete sleep deprivation between experimental day 1 and 2, and 1 was excluded due to medical history. Ultimately, 32 participants (24.16 ± 4.1 years, 18 females, 16 participants per group) completed the study and were included in the final analysis.

The age of the participants ranged from 18 to 40 years. Exclusion criteria included left-handedness, a history of chronic or acute neurological or psychiatric conditions, prior epileptic seizures or a history of epilepsy, pacemaker or deep brain stimulation, current pregnancy, metal implants in the head or neck region, a history of head injury involving skull fractures or brain tissue damage, a history of intracerebral ischemia or bleeding, current use of CNS-acting medication, smoking, alcohol abuse, drug addiction, skin lesions at the site of electrode placement, recent participation in another scientific or clinical study within the past four weeks, previous experience with fear conditioning protocols, sleep between the first and second experimental day, and an absence of fear acquisition.

Participants were recruited through online advertisements and notices displayed on bulletin boards at the campuses of the Technical University Dortmund and Ruhr University Bochum. Handedness was assessed using the Edinburgh Handedness Inventory (Oldfield, 1971). Participants were compensated with 15 euros per hour for their time, with an additional 30 euros provided for the night of sleep deprivation. They were instructed to abstain from alcohol consumption the night before and on the days of the experiment, and to avoid energy drinks and caffeine for at least three hours prior to the sessions. Informed consent was obtained from all participants prior to their involvement in the study. The study received ethical approval from the Leibniz Research Centre for Working Environment and Human Factors and was conducted in accordance with the Declaration of Helsinki. Sample size was determined in accordance with previous research about tDCS effects on fear extinction and SCR as primary outcome measure, i.e. 16 subjects per group (Marković et al., 2021).

### 2.2. Transcranial direct current stimulation

Multi-electrode tDCS was administered using a battery-operated Starstim stimulator (Neuroelectrics, Spain). The current was delivered through five rubber electrodes, each with a 1 cm radius. This configuration included one anodal electrode and four cathodal electrodes, all of which were covered with conductive electrode paste (Ten20, Weaver and Company, USA) and positioned on the scalp. Computational modeling was conducted using ROAST (Huang et al., 2019) to determine the optimal stimulation montage. In the final configuration, the anodal electrode was placed over the nasion, with the four cathodal electrodes positioned at F7, F8, Ex19, and Ex20. These electrode positions were selected to maximize the average electric field strength in the target area, vmPFC, while minimizing the field strength in the dlPFC and ACC, which are also involved in fear and extinction learning (see supplementary material for details).

Real tDCS was applied with a 30-second ramp-up, followed by 15 minutes of stimulation at an intensity of 2 mA, and a 30-second ramp-down. Stimulation duration was selected based on a previous study that enhanced motor memory consolidation during REM sleep (Nitsche et al., 2010). In further alignment with that study protocol, stimulation was administered during the second and, if necessary, the third REM sleep cycle to ensure sufficient REM sleep duration for effective stimulation. The first REM sleep cycle was not used because it is typically too short. To ensure stable REM sleep before initiating stimulation, the procedure began after 1 minute of online REM sleep scoring. Two participants received less than 15 minutes of real tDCS (i.e., 13 and 12 minutes) as they woke up and were unable to fall back asleep.

For sham stimulation, the current was applied with a 30-second ramp-up, followed by 2 mA stimulation for 30 seconds, and then a 30-second ramp-down to 0 mA. Additionally, EMLA cream (Aspen Pharmacare UK Limited, United Kingdom) was applied for all participants to anesthetize the skin under the tDCS electrodes to prevent sensations that might cause awakening.

### 2.3. Fear conditioning protocol

This study employed a 3-day differential fear conditioning paradigm, based on the protocol developed by Milad et al. (2007). The fear acquisition phase took place on the first day and involved participants learning to differentiate between two stimuli: a conditioned stimulus (CS+, danger stimulus) paired with an unconditioned stimulus (US) (i.e., electrical shock) and a conditioned stimulus (CS-, safety stimulus) that was never paired with the US. A total of 32 trials was presented, consisting of 16 CS+ and 16 CS-trials. The US was delivered with a partial reinforcement rate of 62.5%, meaning that the CS+ was paired with the US in 10 out of the 16 trials.

The extinction phase was conducted the following day and involved the presentation of the CS+ without the electrical shock, while the CS-continued not to be paired with the shock. During extinction, a total of 16 trials were presented, consisting of 8 CS+ and 8 CS-trials. The first recall session was conducted on the same day as the extinction phase, while the second recall session took place the following day (i.e., the third day). These recall sessions involved the same task as the extinction phase (see Figure 1).

**Figure 1.**
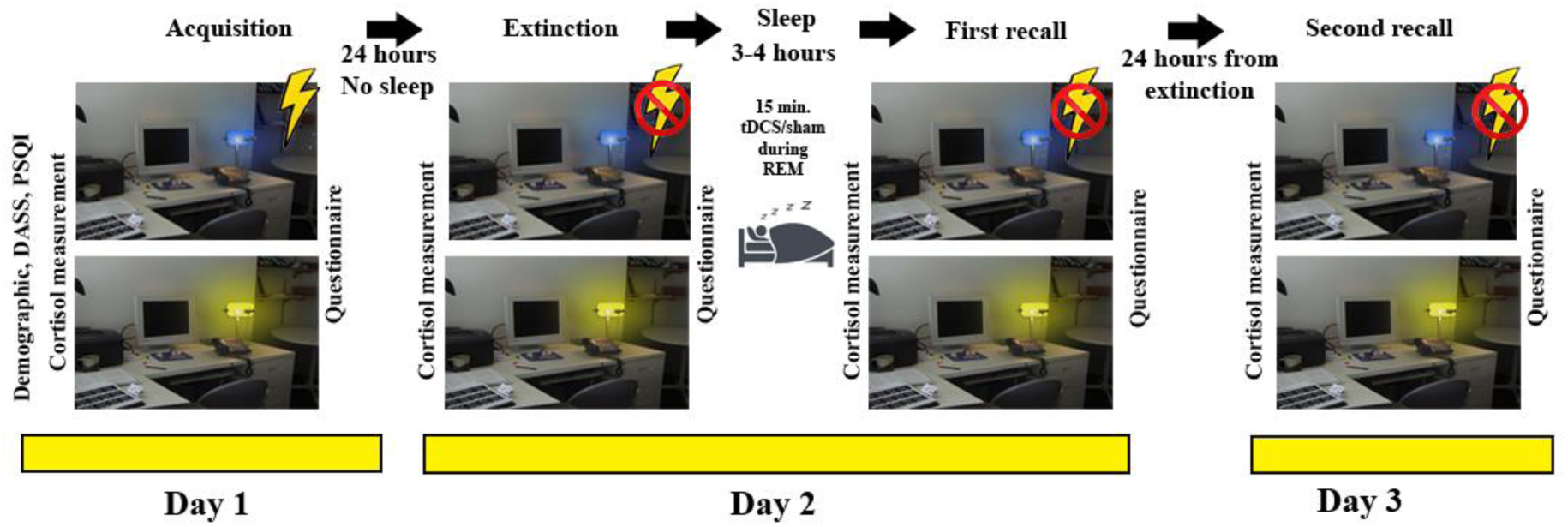
Experimental protocol. The experimental protocol spanned three days, beginning with the fear acquisition phase on Day 1. During this phase, participants learned to differentiate between two stimuli: a blue lamp (CS+), which was paired with an unconditioned stimulus (US), and a yellow lamp (CS-), which was not paired with the US. The acquisition phase lasted for 20 minutes. Between Day 1 and Day 2, participants were sleep-deprived. Approximately 24 hours after fear acquisition, they participated in fear extinction training. In this phase, the CS+ was no longer paired with the US, leading to extinction of the conditioned fear response. The extinction phase lasted for 10 minutes. After extinction training, participants were placed in bed and instructed to sleep, their sleep was monitored via polysomnography. tDCS or sham stimulation was applied during the second REM sleep cycle, and if necessary, during the third cycle, for a total duration of 15 minutes. Following the sleep phase, participants underwent the first recall session, which was identical to the extinction phase. On Day 3, approximately 24 hours after the extinction phase, the second recall session was conducted, following the same procedure as the extinction phase. Cortisol measurements were taken before each phase, and participants completed questionnaires after each phase.

**Figure 2.**
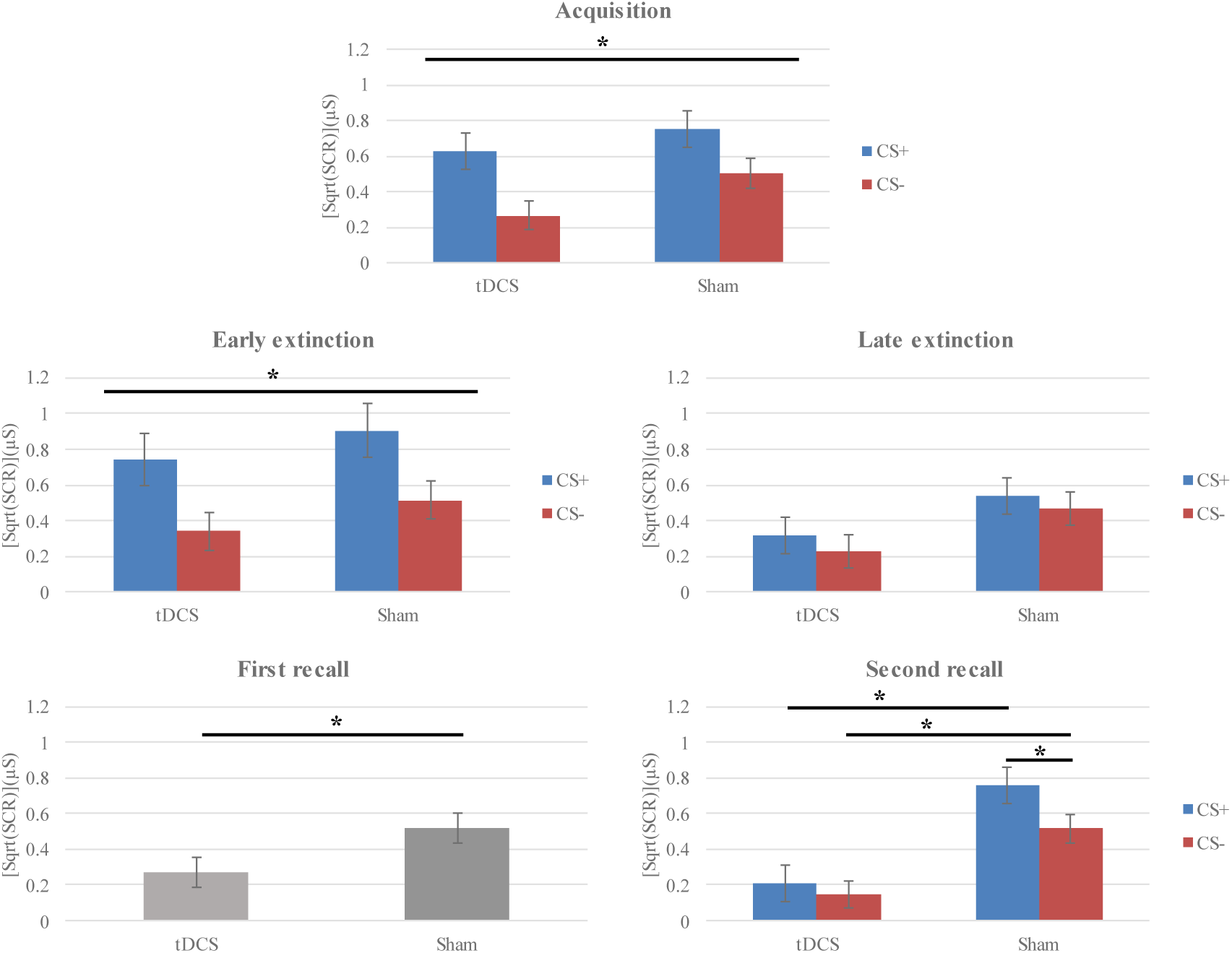
Skin conductance responses during fear acquisition, extinction, first and second recall. *Acquisition*: During the fear acquisition phase, both groups equally acquired fear responses. The horizontal line represents the results of post-hoc tests conducted according to the ‘stimulus’ main effect of the ANOVA in the acquisition phase. *Extinction:* In the early phase of fear extinction, both groups showed fear responses, while the lack of differences during the late extinction phase suggests successful fear extinction in both groups. The horizontal line represents the results of post-hoc tests conducted according to the block × stimulus interaction of the ANOVA for the extinction phase. *First recall:* During the first recall phase, the group which received tDCS showed reduced SCR, irrespective of stimulus condition, compared to sham. The horizontal line represents the results of the post-hoc tests conducted according to the ‘tDCS’ main effect of the ANOVA in first recall phase. *Second recall:* In the second recall phase, the tDCS group showed a lack of differentiation between CS+ and CS-, in contrast to the sham group. Additionally, the tDCS group showed reduced SCR for both CS+ and CS-compared to the sham group, suggesting enhanced fear extinction memory consolidation. The horizontal line represents the results of post-hoc tests conducted according to the tDCS × stimulus interaction of the ANOVA for the second recall phase. * p<0.05; Error bars represent the standard error of the mean. CS+ = conditioned stimulus paired with the unconditioned stimulus; CS-= conditioned stimulus not paired with the unconditioned stimulus; SCR = skin conductance response; sqrt = square root transformation; tDCS = transcranial direct current stimulation; µS = microSiemens.

**Figure 3.**
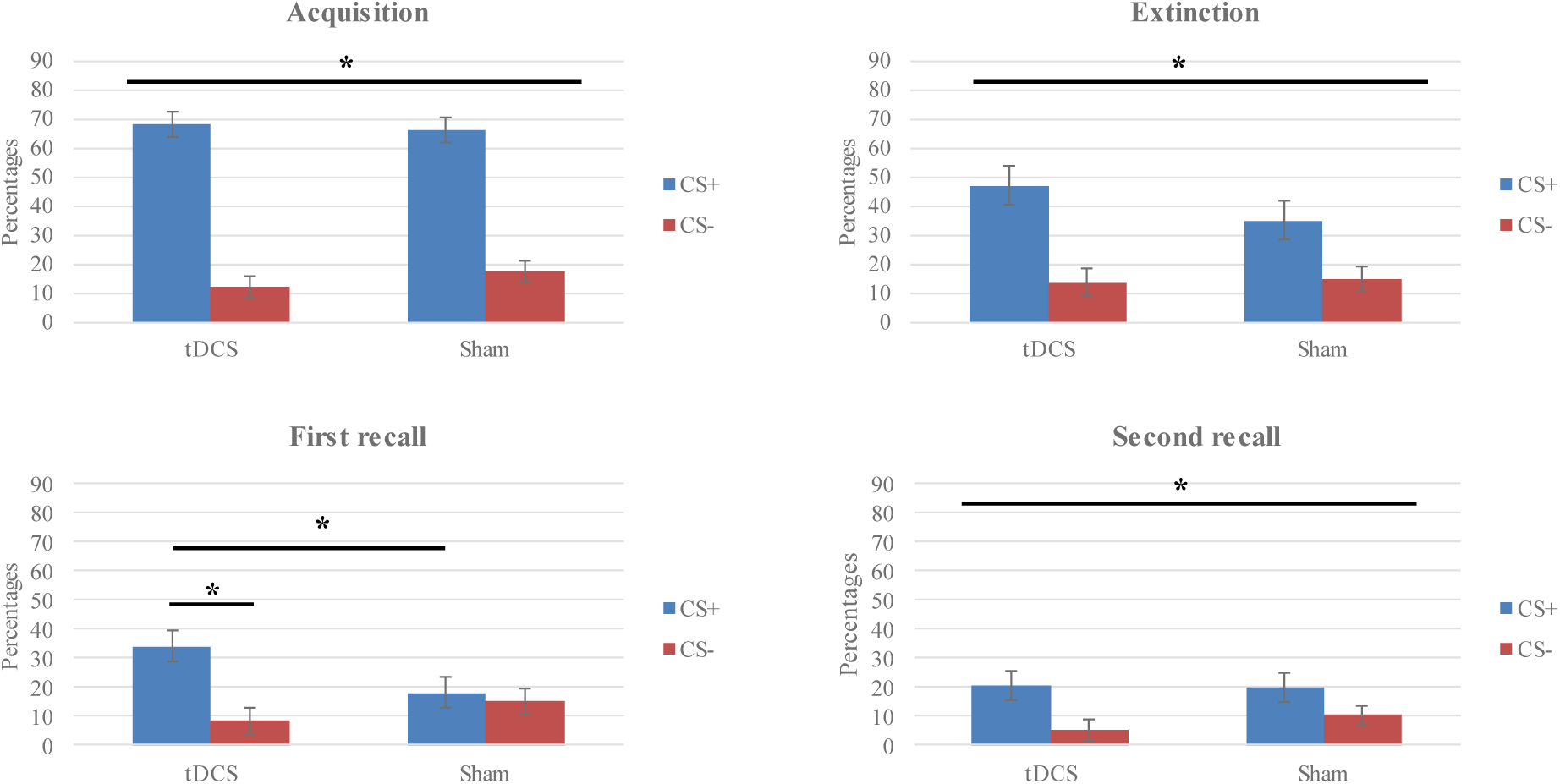
Arousal ratings during fear acquisition, extinction, first and second recall. *Acquisition:* During the fear acquisition phase, both groups successfully differentiated between CS+ and CS-. The horizontal line represents the results of post-hoc tests conducted in accordance with the significant main effect of ‘stimulus’ of the ANOVA in acquisition phase. *Extinction:* In the extinction phase, the differentiation between CS+ and CS-remained for both groups. The horizontal line represents the results of post-hoc tests conducted in accordance with the significant main effect of ‘stimulus’ of the ANOVA in the extinction phase. *First recall:* During the first recall phase, the real tDCS group showed a higher subjective arousal level for the CS+, as compared to the sham group. The horizontal line shows the results of the post-hoc tests conducted in accordance with the significant tDCS × stimulus interaction of the ANOVA in the first recall phase. *Second recall:* In the second recall phase, both groups do not show a differentiation between CS+ and CS-in accordance with the ANOVA results. * p<0.05; Error bars represent standard error of the mean. CS+ = conditioned stimulus paired with unconditioned stimulus; CS-= conditioned stimulus not paired with the unconditioned stimulus; Percentages = marked point on the visual analogue scale transferred to percentages; tDCS = transcranial direct current stimulation.

Each trial lasted for 37.5 seconds. At the beginning of each trial, a fixation cross was presented for a variable duration (0.3 to 1.8 seconds), followed by an image of an office room with a lamp on the desk, displayed for one second. After one second, the lamp in the picture turned either blue or yellow and remained for 12 seconds. Immediately afterward, another fixation cross was shown for a variable duration (22.7 to 24.2 seconds). For the blue lamp condition, the US was applied 100 ms before the end of CS+ (Ma et al., 2024). The US was delivered for the duration of 100 ms and consisted of a short train of four consecutive 500 µs pulses with an inter-pulse interval of 33 ms.

The intensity of the US was calibrated using the method described by Orr et al. (2000). This procedure involved gradually increasing the intensity of electrical shocks and having participants rate the sensation on a scale of 1 to 9, where 1 indicated no or a barely perceived sensation and 9 a painful sensation. The stimulation intensity was set at a level where participants rated the sensation with 8, indicating that it was “highly uncomfortable but not painful.” Stimuli were presented using Presentation software (version 23.0, Neurobehavioral Systems, Inc., USA).

The US was delivered via a ring-shaped electrode placed on the distal phalanx of the right index finger, using a constant current stimulator (DS7A, Digitimer Ltd., UK).

### 2.4. Skin conductance responses

Electrodermal activity (EDA) is a measure of sympathetic nervous system activity and is commonly used in fear conditioning protocols as an indicator of fear responses (Boucsein, 2012; Lonsdorf et al., 2017). EDA was recorded using the Biopac system from two Ag/AgCl electrodes attached to the thenar and hypothenar eminences of the left hand. The electrodes had a contact area with an 11 mm diameter and were covered with isotonic conductive gel.

During SCR recording, a DC amplifier (EDA100C, Biopac Systems, Inc., USA) was employed, and the analog signal was processed using an MP-160 digital converter (Biopac Systems, Inc., USA) at a sampling rate of 1000 Hz. The data were collected in microSiemens (μS) units and stored using AcqKnowledge 5.0 software (Biopac Systems, Inc., USA).

Further data processing and analysis were conducted using MATLAB software (The MathWorks, Inc., United States). Continuous signals were segmented based on the appearance of the CS, with a time window of 1 to 12.5 seconds selected to calculate the SCR. This time window was chosen in accordance with recommendations that the SCR should be derived from data spanning the entire CS-US interval (Pineles et al., 2009). A semi-automated response detection procedure was applied, and the highest trough-to-peak value within the specified time period was identified as the SCR response (Ma et al., 2024). Responses with values less than 0.01 μS were excluded from the analysis (Boucsein, 2012).

### 2.5. Polysomnography

Polysomnographic data were collected using a 64-channel electroencephalography (EEG) system (Bittium NeurOne, Bittium, Finland). The placement of electrodes, visualization of signals, and sleep scoring adhered to the guidelines set by the American Academy of Sleep Medicine (AASM) (Berry et al., 2017). Electrodes were positioned according to the international 10-20 EEG system.

Electrooculogram (EOG) recordings were obtained using two electrodes: one placed 1 cm below and 1 cm lateral to the left outer canthus, and the other positioned 1 cm above and 1 cm lateral to the right outer canthus. The electromyogram (EMG) was recorded with two electrodes attached to the chin, with one electrode positioned 2 cm below the inferior edge of the mandible and 2 cm to the right of the midline, and the other symmetrically placed on the left side. All signals were sampled at a rate of 2000 Hz.

For the purpose of visualizing EEG and EOG data during online sleep scoring, a low-pass filter of 0.3 Hz and a high-pass filter of 35 Hz were applied. For the EMG, the filters were set at 10 Hz for low-pass and 100 Hz for high-pass. During the online scoring process, specific EEG derivations (F4-M1, C4-M1, and O2-M1) were actively monitored, with additional electrodes (F3-M1, C3-M1, and O3-M1) reserved for backup in case of malfunction. EOG and EMG electrodes were displayed in separate channels. Sleep stages were scored in real-time, with 30-second epochs, and participants were continuously monitored via camera and microphone throughout the sleep session.

Offline scoring of the polysomnographic data was conducted manually in Domino software (SOMNOmedics, Germany) following AASM guidelines.

### 2.6. Actigraphy

Actigraphy, a method for monitoring rest and activity cycles, has been widely used to study sleep and wake patterns (Ancoli-Israel et al., 2003). In this study, actigraphy was utilized to ensure that participants adhered to sleep restrictions between the experimental sessions on days 1 and 2. Movement activity was recorded using the MotionWatch 8 device (CamNtech Ltd., England), which participants wore on their wrists. This device provided continuous activity data from the time participants left the laboratory on Day 1 until their return the following day. The recorded activity patterns were visually assessed before the extinction phase, and any participants showing sleep patterns indicated by reduced movement activity were excluded from the experiment.

### 2.7. Cortisol measurement

Stress plays a crucial role in fear learning and extinction processes (Drexler et al., 2019). During stress responses, cortisol levels typically rise, making salivary cortisol a commonly used biomarker for assessing stress (Nicolson, 2008; Bozovic et al., 2013). To monitor and control for stress levels throughout the experiment, saliva samples were collected from participants before each experimental phase (i.e., acquisition, extinction, first recall, and second recall). On the first day, saliva samples were specifically collected prior to adjusting the intensity of the US.

Saliva samples were collected using Salivettes (Sarstedt, Germany). Participants were instructed to place a cotton swab in their mouth for two minutes to ensure a sufficient sample for cortisol analysis. After collection, the saliva samples were stored in a refrigerator for a few days before being processed. The Salivettes were then centrifuged at 1,000 x g for 2 minutes at room temperature. The saliva was aliquoted and stored at −20°C until analysis. Cortisol levels were measured using the Cortisol Saliva ELISA (IBL International, Germany), following the manufacturer’s instructions.

### 2.8. Self-reported measures

Before the experiment, participants completed questionnaires to collect demographic information (i.e., gender, age, weight, height, years of education, intensity of applied electrical shock), the Depression Anxiety Stress Scales (DASS-21; Lovibond & Lovibond, 1995) to ensure there were no significant differences between groups in terms of depression, anxiety, and stress levels, and the Pittsburg sleep quality index (PSQI; Buysse et al., 1989) to assess sleep quality and potential sleep disturbances over a one-month period.

Following each phase of the fear conditioning paradigm (i.e., acquisition, extinction, and first and second recall), participants were asked to complete additional task-related subjective questionnaires, and after the first and second recall a post-stimulation questionnaire to explore if they noticed any side effects of tDCS (Brunoni et al., 2011). Regarding the task-related questionnaire, they were required to respond to questions assessing arousal, fear, and emotional valence of the CS+ and CS-. Arousal and fear levels were rated using a visual analogue scale, with arousal ranging from “calm and relaxed” to “very excited” and fear ranging from “not at all fearful” to “very fearful.” Emotional valence was evaluated on a 5-point Likert scale, ranging from “very unpleasant” to “very pleasant.”

After the acquisition phase, participants were asked additional questions (i.e., acquisition-related questions) to evaluate the effectiveness of fear learning. These included rating the intensity of the last US on a nine-point scale from “not unpleasant” to “very unpleasant,” estimating the number of electrical shocks they had received, specifying the contingency of CS-US (i.e., the percentage of times each CS was followed by an electrical shock), and indicating whether an electrical shock followed each CS (”always,” “sometimes,” “never,” “I don’t know”). Furthermore, participants were asked whether they had recognized a connection between the CS and US, how many electrical shocks it took for them to realize this connection, to decide which color of the lamp was not paired with the electrical shock (by selecting either blue or yellow), and to mark on a visual analogue scale (ranging from the beginning of the task to the end) when they became aware of the CS-US associations during the acquisition phase.

### 2.9. Experimental procedure

Day 1: Acquisition session. The first appointment (acquisition session) began at 11:00 AM. Participants were initially briefed on the details of the experiment and the methods to be used. They were also provided with additional information about the study and asked to sign an informed consent form. A medical doctor then conducted an examination to rule out any potential risks associated with the experiment. Following this, the participants completed a set of questionnaires.

Electrodes for measuring the SCR were attached, and saliva samples were collected to measure cortisol levels. The intensity of the electrical shock was then determined, and participants underwent the fear conditioning task, during which the SCR was recorded. After completing the task, participants filled in the task-related questionnaire (including evaluation of arousal, fear and valence levels for the CS+ and CS-) and acquisition-related questions. Since participants should not sleep until next session, they were given an actigraph to wear until the following day to monitor their activity. This session lasted for approximately 2.5 hours.

Day 2: Extinction session, tDCS/Sham during REM sleep, and first recall session. The second appointment, which also started at 11:00 AM, involved the extinction session, tDCS or sham stimulation during sleep, and the first recall session. Initially, the experimenter assessed the participants’ movement activity over the past 24 hours to ensure they had not slept. If a participant had slept, the experiment was terminated. Otherwise, electrodes for SCR measures were attached, and saliva samples were taken. Participants then underwent the fear extinction procedure. Afterward, they completed the task-related questionnaire and were taken to the sleep laboratory. In the sleep laboratory, tDCS electrodes, along with an EEG cap, and EMG and EOG electrodes, were placed on the participant’s head. Participants were then placed in a dark and soundproof room and instructed to sleep.

During sleep, EEG activity was continuously recorded to monitor sleep stages. tDCS was initiated at the start of REM sleep (i.e., 1 minute after online scoring confirmed the respective sleep stage), with a 30-second ramp-up, followed by 15 minutes of stimulation, and a 30-second ramp-down. Only the second and third REM cycles were considered for stimulation, as the first REM cycle is typically short. Stimulation was initiated during the second REM phase, ramped up for 30 seconds and ended after 15 minutes and ramped down. If REM sleep was shorter, tDCS was paused and continued during the third REM cycle for its remaining predetermined duration. Participants were awakened immediately after stimulation. The sham condition mirrored the real tDCS condition, except that tDCS was ramped up for 30 seconds, applied for 30 seconds, and then ramped down. Participants spent up to 4 hours in bed to ensure sufficient REM sleep for stimulation.

After stimulation, the EEG cap, EMG, EOG, and tDCS electrodes were removed, and participants washed their hair. Before the recall phase, SCR electrodes were reattached, and another saliva sample was taken for cortisol measurement. Approximately 20 to 30 minutes after wakening, participants underwent the first recall phase. After completing task-related and post-stimulation questionnaires, the experimental session concluded. This session lasted approximately 7 hours.

Day 3: Second recall session. The final session (second recall session) took place approximately 24 hours after the fear extinction task. Initially, SCR electrodes were attached, and saliva samples were collected. Participants then underwent the second recall phase and subsequently completed task-related and post-stimulation questionnaires. This session lasted approximately 1 hour. Afterwards, participants were debriefed about the experiment (see Figure 1).

### 2.10. Statistical analysis

A mixed-model ANOVA and ANCOVA were employed to assess group differences in SCR and task-related subjective measures, consistent with previous research (Abend et al., 2016; Vicario et al., 2020; Ney et al. 2021; Ma et al., 2024). These analyses were conducted separately for each phase of the experiment.

The highest SCR value within the predefined time window for each trial was entered as separate observation in the mixed-model ANOVA and ANCOVA as dependent variable. For the fear acquisition phase, an ANOVA was performed with ‘stimulus’ (CS+ or CS-) and ‘trial’ (number of trials, 16 for CS+ and 16 for CS-) as within-subject factors, and ‘tDCS’ (real or sham) as a between-subject factor. For extinction, a separate ANOVA was conducted with ‘block’ (early block (containing the first half of CS+/CS-trials) and late block (containing the second half of CS+/CS-trials) in extinction, first recall, and second recall), ‘stimulus’ (CS+ or CS-), and ‘trial’ (8 trials for CS+ and 8 trials for CS-) as within-subject factors, and ‘tDCS’ as a between-subjects factor. For the first and second recall phases, beyond ANOVAs identical to that described for extinction, separate ANCOVAs were performed, incorporating the same factors as in the extinction phase, and REM sleep duration as a covariate. This adjustment aimed to assess whether REM sleep duration influenced results, given its role in fear extinction memory consolidation (Pace-Schott et al., 2015; Bottary et al., 2023). To normalize the distribution of SCR data, a square root transformation (sqrt) was applied in all conditions.

A similar statistical approach was applied to task-related questionnaire data. Arousal and fear ratings, collected via a visual analogue scale, were converted to percentages for analysis. Valence ratings, initially scored on a 1-5 scale, were reversed so that higher values indicated increased unpleasantness before being entered into the analysis. Separate mixed-model ANOVAs were conducted for arousal, fear, and valence ratings during the acquisition and extinction phases, with ‘stimulus’ (CS+ or CS-) as a within-subject factor and ‘tDCS’ (real or sham) as a between-subjects factor. For the first and second recall phases, separate ANCOVAs were performed with REM sleep duration included as covariate.

To evaluate potential group differences in age, height, weight, education, and intensity of the applied electrical shock, as well as scores on the DASS-21 and PSQI subscales, independent-sample t-tests were conducted, while a chi-square test was performed for assessing gender differences.

Polysomnographic recordings were scored according to AASM guidelines (Berry et al., 2017), with specific parameters extracted, including time in bed (TIB), total sleep time (TST), sleep efficiency, total wakefulness duration, REM sleep duration, N1, N2, and N3 sleep stage durations, along with each sleep stage percentage of total sleep time (%TST). Group differences of sleep parameters were assessed using independent-sample t-tests.

A mixed model ANOVA was used to evaluate differences in cortisol levels measured before each experimental phase. ‘Phase’ (all experimental phases) was included as a within-subject factor, and ‘tDCS’ as between-subject factor.

For all ANOVAs and ANCOVAs, Mauchly’s test of sphericity was applied. If sphericity was violated, the Greenhouse–Geisser correction was implemented. Least significant difference (LSD) post-hoc comparisons were conducted when significant ANOVA or ANCOVA results were found. The critical p-value was set at 0.05 for all statistical analyses, with effect sizes reported as partial eta squared (ηp²). Statistical analyses were conducted using IBM SPSS Statistics version 29.0.

## 3. Results

### 3.1. Demographics

No significant differences were found between the groups regarding gender (χ² = 2.032, df = 1, p = .154), age (t(30) = -.297, p = .768), height (t(30) = −1.618, p = .116), weight (t(30) = -.789, p = .436), or education (t(30) = .680, p = .502). Similarly, no significant differences were observed in depression (t(21.354) = −1.356, p = .189), anxiety (t(30) = .651, p = .520), or stress scores (t(30) = -.140, p = .890) as measured by the DASS-21. Additionally, the intensity of the applied electrical shock did not significantly differ between groups (t(30) = -.981, p = .334). These findings indicate that the experimental groups were comparable in terms of demographic factors, baseline levels of depression, anxiety, and stress, as well as the intensity of the applied electrical shock. No differences were detected regarding the perceived intensity of the applied electrical stimulus (see supplementary material).

Furthermore, no differences between groups were found regarding the global PSQI score (t(29) = .119, p = .906), and its components of subjective sleep quality (t(29) = -.761, p = .453), sleep latency (t(29) = .621, p = .539), sleep duration (t(29) = -.349, p = .730), sleep efficiency(t(29) =.846, p = .405), sleep disturbances (t(15) =1.464, p = .164), use of sleep medications, and daytime dysfunction (t(29) = -.703, p =.488) (Table 1b). One participant did not fill in the PSQI questionnaire.

**Table 1a.**
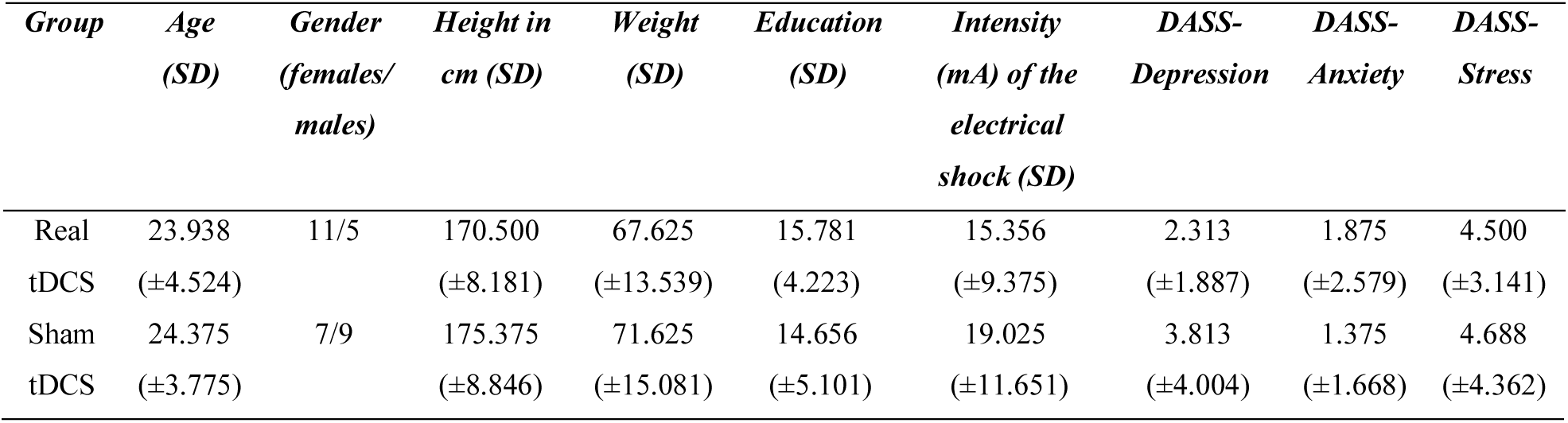
Demographic details, electrical shock intensity and DASS subscales. Values presented are means (M) and standard deviations (SD). DASS-21= Depression, Anxiety, Stress scale; tDCS = transcranial direct current stimulation.

**Table 1b.**
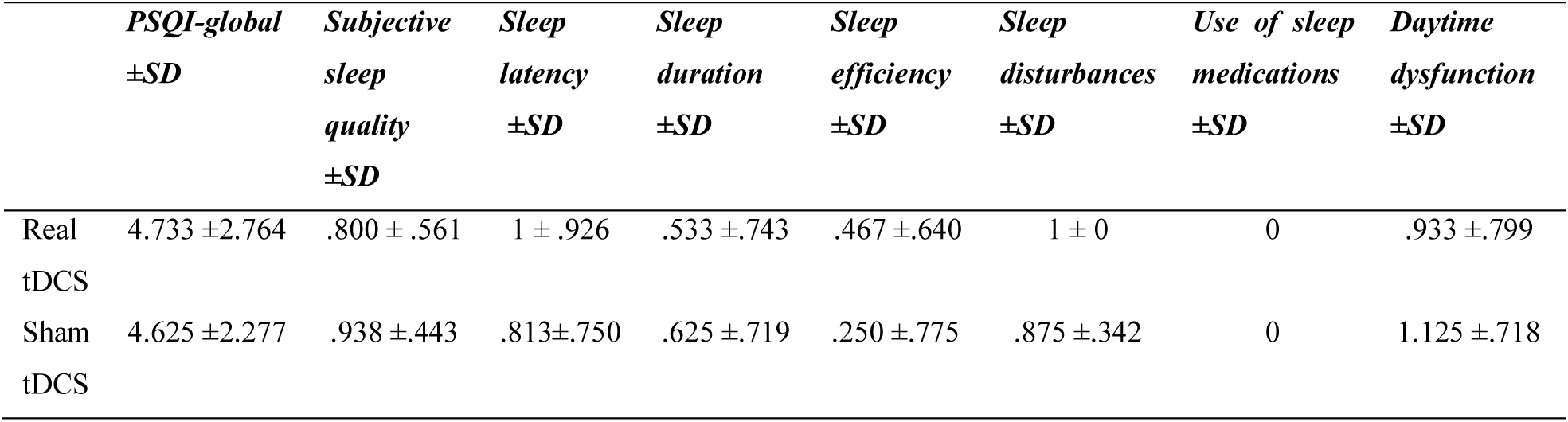
Sleep quality. PSQI = Pittsburg sleep quality index; SD = standard deviation; tDCS = transcranial direct current stimulation.

### 3.2. Cortisol measurement and sleep architecture

No significant main effect of ‘tDCS’ was found (F(1,30) = .523, p = .475). Furthermore, the analysis did not reveal a significant interaction between phase × tDCS (F(2.316, 69.488) = .525, p = .620), indicating that cortisol levels did not significantly differ between groups at any time point. A significant main effect of ‘phase’ (F(2.316, 69.488) = 10.258, p < .001) was however observed.

Significant differences between groups were observed for several sleep parameters. The sham group had increased TIB (t(30) = −2.374, p = .024) and TST (t(30) = −2.548, p = .016) as compared to the real stimulation group. In contrast, the tDCS group exhibited a higher percentage of REM sleep in TST (t(25.947) = 2.322, p = .028). To account for potential confounding effects, an ANCOVA was conducted with REM duration included as a covariate for all outcome measures. The main results remained unchanged. Detailed results of this analysis are provided in the supplementary material. No significant were observed for sleep efficiency (t(30) = -.941, p = .354), total wakefulness duration (t(30) = .476, p = .637), REM sleep duration (t(30) = .883, p = .384), N1 stage duration (t(30) = -.878, p = .378), N1 % TST (t(30) = -.327, p = .746), N2 stage duration (t(30) = −1.361, p = .184), N2 % TST (t(30) = -.133, p = .895), N3 stage duration (t(30) = −1.338, p = .191), N3 % TST (t(30) = -.451, p = .655).

### 3.3. Skin conductance responses

#### Acquisition phase

The ANOVA results show a significant main effect for the factor ‘stimulus’ (F(1,30) = 69.366, p < .001), and exploratory post-hoc tests revealed that SCR for CS+ (M = .691, SEM = .073) were significantly (p < .001) larger than those for CS-(M = .385, SEM = .057). Additionally, a significant main effect was found for the factor ‘trial’ (F(6.928, 207.852) = 4.756, p < .001). No other significant main effects or interactions were identified.

#### Extinction phase

The ANOVA identified significant main effects for ‘block’ (F(1,30) = 19.784, p < .001), ‘trial’ (F(3,30) = 4.637, p = .005), and ‘stimulus’ (F(1,30) = 20.085, p < .001). A significant block × stimulus interaction was also observed (F(1,30) = 21.460, p < .001). No other significant main effects or interactions were found.

Respective post-hoc tests revealed that during early extinction, the SCR for CS+ (M = .823, SEM = .105) was significantly (p < .001) larger than for CS-(M = .428, SEM = .073). However, in the late extinction phase, the difference between CS+ (M = .428, SEM = .069) and CS-(M = .350, SEM = .066) was not statistically significant (p = .138). Additionally, the SCR for CS+ in early extinction was significantly larger than the SCR for CS+ in late extinction (Mdif = .396, p < .001). In contrast, the difference of the SCR for CS-between early and late extinction was not significant (Mdif = .078, p = .082).

#### First recall phase

The ANOVA identified significant main effects for the factors ‘block’ (F(1,30) = 25.290, p < .001), ‘trial’ (F(3,30) = 10.118, p < .001), ‘stimulus’ (F(1,30) = 12.423, p = .001), and ‘tDCS’ (F(1,30) = 4.202, p = .049). Additionally, significant interactions were found for block × trial (F(3,30) = 7.271, p < .001) and block × stimulus (F(1,30) = 5.039, p = .032).

Post-hoc tests based on the main effect of the factor ‘tDCS’ revealed that SCR, irrespective of the stimulus condition, for the group receiving tDCS was significantly lower (p = .049) (M = .273, SEM = .085) compared to sham (M = .521, SEM = .085).

#### Second recall phase

The ANOVA identified significant main effects for factors ‘block’ (F(1,30) = 19.974, p < .001), ‘trial’ (F(2.311, 69.330) = 10.310, p < .001), ‘stimulus’ (F(1,30) = 18.363, p < .001), and ‘tDCS’ (F(1,30) = 13.865, p < .001). Significant interactions were also found for block × stimulus (F(1,30) = 10.749, p = .003), and block × trial (F(3,90) = 3.094, p = .031). Notably, there was a significant tDCS × stimulus interaction (F(1,30) = 6.295, p = .018).

Post-hoc tests based on the tDCS × stimulus interaction revealed that in the sham group, the SCR for CS+ (M = .763, SEM = .102) was significantly larger (p < .001) than for the CS-(M = .517, SEM = .079). In contrast, the real tDCS group showed no significant difference (p = .219) between CS+ (M = .211, SEM = .102) and CS-(M = .147, SEM = .079). Additionally, significant between-group differences of the SCR were observed. The real tDCS group SCR scores were lower for both, the CS+ (Mdif = .552, p < .001) and CS-(Mdif = .370, p = .002) compared to the sham group. No further significant main effects or interactions were identified.

The main results for the first and second recall are consistent across both ANOVAs and ANCOVAs. See the supplementary material for further details.

### 3.4. Tak-related questions

#### 3.4.1. Arousal ratings

##### Acquisition

The ANOVA revealed a significant main effect of the factor ‘stimulus’ (F(1,30) = 158.462, p < .001), and respective post-hoc tests revealed that arousal ratings for the CS+ (M = 67.028, SEM = 3.117) were significantly (p < .001) higher than those for the CS-(M = 14.839, SEM = 2.583). No further significant main effects or interactions were identified. *Extinction:* The ANOVA revealed a significant main effect of the factor ‘stimulus’ (F(1,30) = 39.841, p < .001). Post-hoc tests revealed that ratings for the CS+ (M = 40.999, SEM = 4.689) were significantly (p < .001) higher than for CS-(M = 14.334, SEM = 3.097). No further significant main effects or interactions were identified. *First recall:* The ANOVA identified a significant main effect of the factor ‘stimulus’ (F(1,20) = 17.949, p < .001), along with a significant group × stimulus interaction (F(1,30) = 11.442, p = .002). Post-hoc tests indicated that the tDCS group reported significantly higher arousal ratings for CS+ (M = 34.207, SEM = 5.117) compared to CS-(M = 8.496, SEM = 4.529) (p < .001). In contrast, the sham group showed no significant difference between CS+ (M = 18.189, SEM = 5.117) and CS-(M = 15.307, SEM = 4.529) (p = .550). Additionally, the arousal rating for CS+ in the tDCS group was significantly higher than for CS+ in the sham group (Mdif = 16.018, p = .035), while no significant differences were found for CS-(Mdif = −6.811, p = .296). No further significant main effects or interactions were identified. *Second recall:* The ANOVA detected a significant main effect of the factor ‘stimulus’ (F(1,30) = 24.525, p < .001). Arousal ratings for CS+ (M = 20.228, SEM = 3.558) were significantly higher than for CS-(M = 7.784, SEM = 2.542). No further significant main effects or interactions were identified.

#### 3.4.2. Fear ratings

##### Acquisition

The ANOVA revealed a significant main effect of the factor ‘stimulus’ (F(1,30) = 101.350, p < .001), and respective post-hoc tests revealed that fear ratings for the CS+ (M = 56.905, SEM = 4.473) were significantly (p < .001) higher than for the CS-(M = 7.953, SEM = 1.603). No further significant main effects or interactions were identified. *Extinction*: The ANOVA showed a significant main effect of the factor ‘stimulus’ (F(1,30) = 29.658, p < .001). Post-hoc tests revealed that fear ratings for the CS+ (M = 35.740, SEM = 5.336) were significantly higher (p < .001) than for the CS-(M = 9.618, SEM = 2.711). No further significant main effects or interactions were identified. *First recall:* The ANOVA identified a significant main effect of the ‘stimulus’ factor (F(1,30) = 18.539, p < .001), with CS+ ratings (M = 24.439, SEM = 4.019) significantly higher than CS-ratings (M = 8.046, SEM = 2.255). No further significant main effects or interactions were identified. *Second recall:* The ANOVA revealed a significant main effect of the ‘stimulus’ factor (F(1,30) = 16.549, p < .001). Fear ratings for CS+ (M = 15.924, SEM = 2.995) were significantly higher than for CS-(M = 4.978, SEM = 1.651). No further significant main effects or interactions were identified.

#### 3.4.3. Valence ratings

##### Acquisition

The ANOVA revealed a significant main effect of the factor ‘stimulus’ (F(1,30) = 110.205, p < .001), and respective post-tests revealed that valence ratings for the CS+ (M = 3.750, SEM = .116) were significantly (p < .001) higher than for the CS-(M = 1.656, SEM = .140). No further significant main effects or interactions were identified. *Extinction:* The ANOVA detected a significant main effect of the factor ‘stimulus’ (F(1,30) = 26.193, p < .001). Post-hoc tests revealed that valence ratings for the CS+ (M = 2.969, SEM = .171) were significantly higher (p < .001) than for the CS-(M = 1.906, SEM = .147). No further significant main effects or interactions were identified. *First recall:* The ANOVA identified a significant main effect of the ‘stimulus’ factor (F(1,30) = 19.982, p < .001), with CS+ ratings (M = 2.406, SEM = .148) being significantly higher than CS-ratings (M = 1.813, SEM = .140). No further significant main effects or interactions were identified. *Second recall:* The ANOVA revealed a significant main effect of the ‘stimulus’ factor (F(1,30) = 13.992, p < .001). Valence ratings for CS+ (M = 2.375, SEM = .168) were significantly higher than for CS-(M = 1.781, SEM = .149). No further significant main effects or interactions were identified.

The main results of task-related questions for the first and second recall are consistent across both ANOVAs and ANCOVAs. See the supplementary material for further details.

Answers to the acquisition-related questions (i.e., estimation of the number of US received, contingency of CS-US, recognition of the connection between the CS and US, and when this has occurred) suggested identical patterns of successful fear learning in both groups. Furthermore, blinding was successfully maintained throughout the experiment and no group differences emerged in tDCS-related side effects (for details see Supplementary material).

## 4. Discussion

Previous studies have highlighted the role of REM sleep in emotional regulation and fear extinction memory consolidation (Genzel et al., 2015; Pace-Schott et al., 2015; Bottary et al., 2023). However, current findings are largely correlative, and none have causally assessed the role of REM sleep in fear extinction memory using tDCS. Our findings demonstrate that anodal tDCS applied over the vmPFC during REM sleep enhances fear extinction memory consolidation, as evidenced by reduced SCR during the first and second recall. However, subjective arousal ratings during the first recall indicated that the tDCS group experienced heightened arousal for the CS+.

Both groups successfully acquired and extinguished fear responses as measured by SCR. An immediate effect of tDCS on fear extinction memory consolidation was observed during the first recall. This aligns with the results of a study showing enhancement of motor memory consolidation immediately following tDCS applied during REM sleep (Nitsche et al., 2010). An extended beneficial effect of tDCS on fear extinction memory consolidation was moreover observed during the second recall. However, differences in the observed results between the first and second recall are noteworthy. During the first recall, the reduction in SCR across all trials, regardless of stimulus condition, suggests that tDCS initially modulates general physiological arousal rather than discrimination processes (Lonsdorf et al., 2017). Given that memory consolidation is time-dependent, the full effects of tDCS may take longer to emerge. This is supported by results from the second recall on the following day, where a further reduction in SCR to both CS+ and CS-in the tDCS group, along with a lack of discrimination, indicates enhanced fear extinction consolidation.

REM sleep has been previously shown to be critical for fear extinction memory consolidation in both animals (Silvestri, 2005; Fu et al., 2007; Jung & Noh, 2021) and humans (Spoormaker et al., 2010, 2012; Menz et al., 2016; Bottary et al., 2023; Friesen et al., 2022). The results of the present study align with those findings. Mechanistically, the vmPFC/IL, which was upregulated in the present study by anodal tDCS, has been consistently shown to be involved in fear extinction, with a proposed mechanism involving top-down inhibition of the amygdala, leading to reduced fear responses (Milad et al., 2007; Milad & Quirk, 2012). Furthermore, inducing LTP-like plasticity in the vmPFC via tDCS has been shown to enhance extinction (Marković et al., 2021; Adams et al., 2023; Boehme et al., 2024) and the role of vmPFC regulation of the amygdala during REM sleep has also been emphasized (Walker & van Der Helm, 2009; Spoormaker et al., 2010). This proposed mechanism of action is further supported by a recent optogenetic study showing that suppressing the vmPFC/IL during REM sleep, in contrast to SWS, after fear learning leads to deficits in fear extinction recall (Hong et al., 2024). Accordingly, the present findings suggest that tDCS-induced LTP-like plasticity in the vmPFC enhances the consolidation of fear extinction memory during REM sleep, likely by strengthening top-down regulation of the amygdala. This enhanced consolidation ultimately results in a more effective reduction of physiological fear responses. Overall, the findings of the present study align with the “sleep to remember, sleep to forget” model, which posits that a crucial function of REM sleep is to reduce vegetative reactions induced by emotionally charged experiences in order to enhance adaptive behavior (Walker & van Der Helm, 2009).

The subjective ratings have shown increased arousal ratings after stimulation, which may indicate a potential deficit in fear extinction. This finding requires further consideration. Since only arousal levels were affected, not fear or valence ratings, this may reflect an increase in the subjective importance of the stimuli rather than heightened negative emotions (Kuppens et al, 2013). Alternatively, as previously shown, tDCS might improve arousal and vigilance levels, as measured by behavioral paradigms (McIntire et al., 2017, 2020), which might not be fully captured by SCR measurements. However, this isolated finding warrants further exploration and replication.

Although task-related subjective measures indicated successful fear learning in the present study, they generally did not show a tDCS effect, consistent with previous studies (Dittert et al., 2018; Boehme et al., 2024; Ma et al., 2024). One reason could be that subjective measures, typically assessed at the end of the task, capture general impressions and may not detect subtle differences revealed by trial-based SCR data (Grillon et al., 2007). Additionally, different neural and cognitive processes may underlie more conscious evaluations of stimuli and psychophysiological responses like SCR (LeDoux & Pine, 2016) and the proposed tDCS effect on vmPFC-amygdala connections might be limited to physiological responses.

In the past decade, there has been growing interest in the application of non-invasive brain stimulation methods to treat anxiety disorders, with promising results (Vicario et al., 2019; Jafari et al., 2021; Hui et al., 2024). These approaches could enhance the efficacy of exposure therapy, the gold standard for treating ADs and PTSD (Craske et al., 2014). The results of the present study align with this direction. Applying tDCS over the vmPFC during REM sleep after exposure sessions could potentially enhance the consolidation of extinguished memories and improve therapeutic outcomes.

Some limitations of this study should be considered. Sleep deprivation can impair fear memory consolidation in both rodents (Kumar et al., 2012; Montes-Rodríguez et al., 2019) and humans, leading to reduced recall of fear memory after acquisition and extinction (Menz et al., 2013, 2016). As a result, sleep deprivation may decrease fear memory consolidation and lead to poorer memory strength, which is more quickly extinguished. Hormonal fluctuations, particularly cortisol, which vary between day and night and are heightened after sleep deprivation, could also influence learning outcomes (Wright et al., 2015; Drexler et al., 2019). Therefore, replications with full-night sleep are necessary. Additionally, slow-wave sleep (SWS) might play a role in fear extinction memory consolidation (Hauner et al., 2013; He et al., 2015), suggesting that targeting SWS with tDCS could help to differentiate the roles of REM and SWS in this process. Furthermore, the question of how long the effect of tDCS will last and whether it will be transferable to individuals with ADs needs to be resolved in future studies. Finally, comparing this approach to the standard method of applying stimulation while awake is essential to assess whether targeting REM sleep leads to better fear extinction memory consolidation.

### Conclusion

This study provides evidence that anodal tDCS over the vmPFC during REM sleep enhances fear extinction memory consolidation. While increased subjective arousal post-stimulation requires further investigation, the SCR results support the potential of tDCS during REM sleep to improve therapeutic outcomes in ADs and PTSD. Future studies should consider the influence of sleep deprivation and hormonal fluctuations on the results of the study, and explore the differential roles of REM and SWS in fear extinction memory consolidation.

## Author Contributions

Vuk Marković contributed to design of the study, data collection, data analysis and visualization, writing the early version of the manuscript, reviewing and editing. Gaetano Rizzo contributed to data collection and manuscript reviewing and editing. Fatemeh Yavari contributed to manuscript reviewing and editing. Carmelo M. Vicario and Michael A. Nitsche conceived the work, designed the study and contributed to manuscript reviewing and editing. All authors revised and approved the final version of the manuscript.

## Funding

This work was supported by BIAL foundation (Prot. Number 160/18) and the Deutsche Forschungsgemeinschaft (DFG, German Research Foundation)-Projektnummer 316803389 - SFB 1280, subproject A06. Vuk Marković work was supported from Deutscher Akademischer Austauschdienst (DAAD).

## Competing interests

Michael A. Nitsche is a member of Scientific Advisory Boards of Neuroelectrics and Precisis. The other authors declare no potential conflicts of interest.

## Data Availability

The datasets generated and/or analyzed during the current study are not publicly available due to institutional regulations on data protection and the informed consent provided by participants, but further inquiries can be directed to the corresponding author/s.

