## Supplementary material for "Plasticity induction in the ventromedial prefrontal cortex during REM sleep improves fear extinction memory consolidation": suppl

#### 1. Computational modeling

A computational modeling approach using ROAST (Huang et al., 2019) was utilized to determine electrode positions for tDCS. The criteria were to find the position of the electrodes that will lead to the highest electrical field in the vmPFC, but no relevant electrical field in other brain regions that are relevant for fear expression, such as the dlPFC, ACC, and the cerebellum. The final montage included five electrodes with 1 cm radius; the anode was placed over the nasion while 4 cathodal electrodes were placed over F7, F8, Ex19, and Ex20 electrode positions according to the international 10-20 system (see Figure 1).

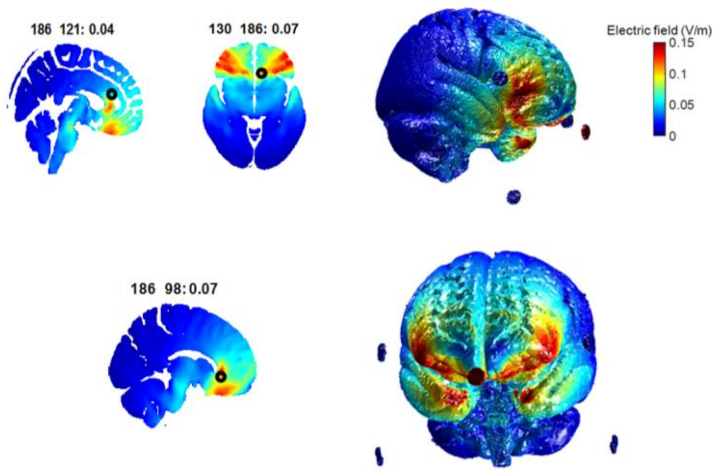


Figure 1. Electrical field distribution for the modeled montage to induce the highest electrical field in the vmPFC.

#### 2. Mixed model ANOVA and ANCOVA results: skin conductance responses

| *Acquisition* | *df* | *F* | *Sig.* | $\eta_p^2$ |
|---|---|---|---|---|
| **stimulus** | 1, 30 | 69.366 | **<.001** | .698 |
| **trial** | 6.928,207.852 | 4.756 | **<.001** | .137 |
| tDCS | 1,30 | 2.016 | .166 | .063 |

| | | | | |
|---|---|---|---|---|
| stimulus × tDCS | 1, 30 | 2.389 | .133 | .074 |
| trial × tDCS | 6.928,207.852 | 1.202 | .303 | .039 |
| stimulus × trial | 8.096, 242.872 | 1.245 | .273 | .040 |
| stimulus × trial × tDCS | 8.096, 242.872 | .799 | .606 | .026 |

Table 1. ANOVA results for the acquisition phase. df = degrees of freedom; F-statistic = calculated statistic for ANOVA; Sig. = significance level (p-value); tDCS = transcranial direct current stimulation; $\eta_p^2$ = Partial Eta Squared (effect size index).

| ***Extinction*** | ***df*** | ***F*** | ***Sig.*** | $\boldsymbol{\eta_p^2}$ |
|---|---|---|---|---|
| **block** | 1,30 | 19.784 | **<.001** | .397 |
| **stimulus** | 1,30 | 20.085 | **<.001** | .401 |
| **trial** | 3,30 | 4.637 | **.005** | .134 |
| tDCS | 1,30 | 2.184 | .150 | .068 |
| **block** × **stimulus** | 1,30 | 21.460 | **<.001** | .417 |
| block × trial | 3,90 | 1.619 | .190 | .051 |
| block × tDCS | 1,30 | .338 | .565 | .011 |
| stimulus × trial | 3,90 | 1.095 | .355 | .035 |
| stimulus × tDCS | 1,30 | .017 | .896 | .001 |
| trial × tDCS | 3,30 | 1.506 | .218 | .048 |
| block × stimulus × trial | 3,90 | 1.820 | .149 | .057 |
| block × stimulus × tDCS | 1,30 | .000 | .998 | .000 |
| block × trial × tDCS | 3,90 | 1.890 | .137 | .059 |
| stimulus × trial × tDCS | 3,90 | .502 | .682 | .016 |
| block × stimulus × trial × tDCS | 3,90 | .880 | .454 | .029 |

Table 2. ANOVA results for the extinction phase. df = degrees of freedom; F-statistic = calculated statistic for ANOVA; Sig. = statistical significance (p-value); tDCS = transcranial direct current stimulation; $\eta_p^2$ = Partial Eta Squared (effect size index).

| ***First recall*** | ***df*** | ***F*** | ***Sig.*** | $\boldsymbol{\eta_p^2}$ |
|---|---|---|---|---|
| **block** | 1,30 | 25.290 | **<.001** | .457 |
| **stimulus** | 1,30 | 12.423 | **.001** | .293 |
| **trial** | 3,30 | 10.118 | **<.001** | .252 |
| **tDCS** | 1,30 | 4.202 | **.049** | .123 |
| **block** × **stimulus** | 1,30 | 5.039 | **.032** | .144 |
| **block** × **trial** | 3,90 | 7.271 | **<.001** | .195 |
| block × tDCS | 1,30 | .862 | .361 | .028 |
| stimulus × trial | 3,90 | 2.025 | .116 | .063 |
| stimulus × tDCS | 1,30 | .020 | .889 | .001 |
| trial × tDCS | 3,30 | 2.057 | .112 | .064 |
| block × stimulus × trial | 3,90 | .846 | .472 | .027 |
| block × stimulus × tDCS | 1,30 | .257 | .616 | .008 |
| block × trial × tDCS | 3,90 | .046 | .987 | .002 |
| stimulus × trial × tDCS | 3,90 | .627 | .599 | .020 |
| block × stimulus × trial × tDCS | 3,90 | .164 | .920 | .005 |

Table 3. ANOVA results for the first recall phase. df = degrees of freedom df = degrees of freedom; F-statistic = calculated statistic for ANOVA; Sig. = statistical significance (p-value); tDCS = transcranial direct current stimulation; $\eta_p^2$ = Partial Eta Squared (effect size index).

| ***First recall*** | ***df*** | ***F*** | ***Sig.*** | $\eta_p^2$ |
|---|---|---|---|---|
| block | 1,29 | .040 | .842 | .001 |
| stimulus | 1,29 | 1.566 | .221 | .051 |
| trial | 3,87 | .386 | .763 | .013 |
| REM | 1,29 | .487 | .491 | .017 |
| **tDCS** | 1,29 | 4.484 | **.043** | .134 |
| block × stimulus | 1,29 | .135 | .716 | .005 |
| block × trial | 3,87 | .514 | .647 | .017 |
| block × REM | 1,29 | 1.567 | .221 | .051 |
| block × tDCS | 1,29 | 1.264 | .270 | .042 |
| stimulus × trial | 3,87 | .672 | .572 | .023 |
| **stimulus × REM** | 1,29 | 5.728 | **.023** | .165 |
| stimulus × tDCS | 1,29 | .281 | .600 | .010 |
| trial × REM | 3,87 | 1.255 | .295 | .041 |
| trial × tDCS | 3,87 | 1.631 | .188 | .053 |
| block × stimulus × trial | 3,87 | .450 | .718 | .015 |
| block × stimulus × REM | 1,29 | .064 | .802 | .002 |
| block × stimulus × tDCS | 1,29 | .284 | .598 | .010 |
| block × trial × REM | 3,87 | .005 | 1 | .000 |
| block × trial × tDCS | 3,87 | .044 | .988 | .002 |
| stimulus × trial × REM | 3,87 | 1.246 | .298 | .041 |
| stimulus × trial × tDCS | 3,87 | .483 | .695 | .016 |
| block × stimulus × trial × REM | 3,87 | .714 | .546 | .024 |
| block × stimulus × trial × tDCS | 3,87 | .115 | .951 | .004 |

Table 4. ANCOVA results for the first recall phase. df = degrees of freedom; F-statistic = calculated statistic for ANCOVA; Sig. = statistical significance (p-value); tDCS = transcranial direct current stimulation; $\eta_p^2$ = Partial Eta Squared (effect size index).

| ***Second recall*** | ***df*** | ***F*** | ***Sig.*** | $\eta_p^2$ |
|---|---|---|---|---|
| **block** | 1,30 | 19.974 | **<.001** | .400 |
| **stimulus** | 1,30 | 18.363 | **<.001** | .380 |
| **trial** | 2.311,69.330 | 10.310 | **<.001** | .256 |
| **tDCS** | 1,30 | 13.865 | **.001** | .316 |
| **block** × **stimulus** | 1,30 | 10.749 | **.003** | .264 |
| **block** × **trial** | 3,90 | 3.094 | **.031** | .093 |
| block × tDCS | 1,30 | 3.420 | .074 | .102 |
| stimulus × trial | 3,90 | .507 | .678 | .017 |
| **stimulus** × **tDCS** | 1,30 | 6.295 | **.018** | .173 |

| | | | | |
|---|---|---|---|---|
| trial × tDCS | 2.311,69.330 | .285 | .784 | .009 |
| block × stimulus × trial | 3,90 | .471 | .703 | .015 |
| block × stimulus × tDCS | 1,30 | .448 | .508 | .015 |
| block × trial × tDCS | 3,90 | .841 | .475 | .027 |
| stimulus × trial × tDCS | 3,90 | .543 | .654 | .018 |
| block × stimulus × trial × tDCS | 3,90 | .142 | .935 | .005 |

Table 5. ANOVA results for the second recall phase. df = degrees of freedom; F-statistic = calculated statistic for ANOVA; Sig. = statistical significance (p-value); tDCS = transcranial direct current stimulation; $\eta p^2$ = Partial Eta Squared (effect size index).

| ***Second recall*** | ***df*** | ***F*** | ***Sig.*** | ***$\eta_p^2$*** |
|---|---|---|---|---|
| block | 1,29 | .939 | .341 | .031 |
| stimulus | 1,29 | .067 | .797 | .002 |
| **trial** | 2.257,65.454 | 3.356 | **.036** | .104 |
| REM | 1,29 | .067 | .798 | .002 |
| **tDCS** | 1,29 | 13.393 | **.001** | .316 |
| block × stimulus | 1,29 | .437 | .514 | .015 |
| block × trial | 3,87 | 2.603 | .057 | .082 |
| block × REM | 1,29 | .066 | .799 | .002 |
| block × tDCS | 1,29 | 3.378 | .076 | .104 |
| stimulus × trial | 3,87 | .853 | .469 | .029 |
| stimulus × REM | 1,29 | .924 | .344 | .031 |
| **stimulus** × **tDCS** | 1,29 | 6.9 | **.014** | .192 |
| trial × REM | 2.257,65.454 | 1.187 | .315 | .039 |
| trial × tDCS | 2.257,65.454 | .149 | .885 | .005 |
| block × stimulus × trial | 3,87 | .907 | .441 | .03 |
| block × stimulus × REM | 1,29 | .058 | .812 | .002 |
| block × stimulus × tDCS | 1,29 | .474 | .497 | .016 |
| block × trial × REM | 3,87 | 1.371 | .257 | .045 |
| block × trial × tDCS | 3,87 | .595 | .62 | .02 |
| stimulus × trial × REM | 3,87 | .656 | .581 | .022 |
| stimulus × trial × tDCS | 3,87 | .594 | .621 | .02 |
| block × stimulus × trial × REM | 3,87 | .638 | .592 | .022 |
| block × stimulus × trial × tDCS | 3,87 | .127 | .944 | .004 |

Table 6. ANCOVA results for the second recall phase. df = degrees of freedom; F-statistic = calculated statistic for ANCOVA; Sig. = statistical significance (p-value); tDCS = transcranial direct current stimulation; $\eta_p^2$ = Partial Eta Squared (effect size index).

### 3. Mixed model ANOVA and ANCOVA results: arousal, fear and valence ratings

| ***Arousal ratings*** | ***Acquisition*** | | | | ***Extinction*** | | | ***First recall*** | | | ***Second recall*** | | |
|---|---|---|---|---|---|---|---|---|---|---|---|---|---|
| | df | F | Sig | $\eta_p^2$ | F | Sig | $\eta_p^2$ | F | Sig. | $\eta_p^2$ | F | Sig. | $\eta_p^2$ |

| | | | | | | | | | | | | | |
|---|---|---|---|---|---|---|---|---|---|---|---|---|---|
| **stimulus** | 1,30 | 158.462 | **<.001** | .841 | 39.841 | **<.001** | .570 | 17.949 | **<.001** | .374 | 24.525 | **<.001** | .450 |
| tDCS | 1,30 | .140 | .711 | .005 | .636 | .431 | .021 | .600 | .445 | .020 | .188 | .668 | .006 |
| **stimulus × tDCS** | 1,30 | .682 | .415 | .022 | 2.362 | .135 | .073 | 11.442 | **.002** | .276 | 1.166 | .289 | .037 |
| ***Fear ratings*** | | | | | | | | | | | | | |
| **stimulus** | 1,30 | 101.350 | **<.001** | .772 | 29.658 | **<.001** | .497 | 18.539 | **<.001** | .382 | 16.549 | **<.001** | .356 |
| tDCS | 1,30 | .817 | .373 | .027 | .056 | .815 | .002 | .481 | .493 | .016 | 1.059 | .312 | .034 |
| stimulus × tDCS | 1,30 | .026 | .873 | .001 | .020 | .889 | .001 | .891 | .353 | .029 | .011 | .918 | .000 |
| ***Valence ratings*** | | | | | | | | | | | | | |
| **stimulus** | 1,30 | 110.205 | **<.001** | .786 | 26.193 | **<.001** | .466 | 19.982 | **<.001** | .400 | 13.992 | **<.001** | .318 |
| tDCS | 1,30 | .332 | .569 | .011 | .599 | .445 | .020 | .374 | .546 | .012 | .013 | .910 | .000 |
| stimulus × tDCS | 1,30 | 1.989 | .169 | .062 | .363 | .552 | .012 | 1.384 | .249 | .044 | 1.899 | .178 | .060 |

Table 7. ANOVA results for arousal, fear and valence ratings in all phases of the experiment. df = degrees of freedom; F-statistic = calculated statistic for ANOVA; Sig. = statistical significance (p-value); tDCS = transcranial direct current stimulation; ηp² = Partial Eta Squared (effect size index).

| | | ***First recall*** | | | ***Second recall*** | | |
|---|---|---|---|---|---|---|---|
| ***Arousal*** | df | F | Sig. | $\eta_p^2$ | F | Sig. | $\eta_p^2$ |
| stimulus | 1,29 | .125 | .726 | .004 | 1.362 | .253 | .045 |
| tDCS | 1,29 | .791 | .318 | .027 | .029 | .867 | .001 |
| REM | 1,29 | .658 | .424 | .022 | 2.94 | .097 | .092 |
| stimulus × REM | 1,29 | 2.622 | .116 | .083 | .035 | .853 | .001 |
| stimulus × tDCS | 1,29 | 10.054 | **.004** | .257 | 1.038 | .317 | .035 |
| ***Fear*** | | | | | | | |
| stimulus | 1,29 | 1.585 | .218 | .052 | 1.853 | .184 | .06 |
| tDCS | 1,29 | .62 | .438 | .021 | .652 | .426 | .022 |
| REM | 1,29 | .466 | .500 | .016 | 1.995 | .168 | .064 |
| stimulus × REM | 1,29 | .008 | .928 | .000 | .070 | .794 | .002 |
| stimulus × tDCS | 1,29 | .866 | .36 | .029 | .021 | .887 | .001 |
| ***Valence*** | | | | | | | |
| stimulus | 1,29 | .019 | .891 | .001 | .001 | .972 | .000 |
| tDCS | 1,29 | .572 | .456 | .019 | .077 | .784 | .003 |
| REM | 1,29 | .928 | .343 | .031 | 1.066 | .310 | .035 |
| stimulus × REM | 1,29 | 1.321 | .260 | .044 | 1.085 | .306 | .036 |
| stimulus × tDCS | 1,29 | .969 | .333 | .032 | 1.432 | .241 | .047 |

Table 8. ANCOVA results for arousal, fear and valence ratings in first and second recall phases. df = degrees of freedom; F-statistic = calculated statistic for ANCOVA; Sig. = statistical significance (p-value); tDCS = transcranial direct current stimulation; ηp² = Partial Eta Squared (effect size index).

### 4. Acquisition-related questions

Independent sample t-tests were conducted to assess group differences in responses to acquisition-related questions. All participants confirmed receiving an electric shock, with the intensity of the last shock rated at an average of 7.77 (SD ± 0.75). There were no significant differences between the groups in shock intensity ratings ($t(30) = -0.116$, $p = .909$). Participants reported receiving an average of 10.53 (SD ± 3.35) shocks, which closely matched the actual number of 10 shocks administered, with no significant group differences found ($t(30) = 1.332$, $p = .193$).

Regarding participants' estimates of the likelihood of receiving a shock after the presentation of the blue or yellow lamp, responses were as follows: for the blue lamp, participants either reported "every time" (53.1%) or "sometimes" (46.9%), while for the yellow lamp, the vast majority (96.9%) chose "never," with only one participant selecting "sometimes" (3.1%). Participants estimated the probability of receiving a shock after the blue lamp at 63.84% (SD ± 15.46), which closely approximated the actual probability of 62.5%. No significant group differences were found in these estimates ($t(30) = -0.877$, $p = .387$). After presentation of the yellow lamp, 31 participants reported a 0% probability of receiving a shock, with one participant estimating a 10% probability.

All participants, except for one who was uncertain, recognized the association between the lamp color and the shock. This occurred early in the acquisition phase, with participants marking an average of 18.2% (SD ± 14.69) on the visual analogue scale, showing no significant group differences ($t(30) = -0.156$, $p = .877$). Participants generally recognized the relationship after experiencing an average of 3 shocks, again with no significant differences between the groups ($t(30) = 0.094$, $p = .926$). When asked to identify which lamp color was not followed by a shock, all participants correctly indicated the yellow lamp.

### 5. tDCS side effects and blinding assessment

A chi-square test was conducted to evaluate the effectiveness of blinding in the tDCS application. The results showed no significant differences between the groups ($\chi^2 = .183$, $df = 1$, $p = .669$), indicating that blinding was successfully maintained throughout the experiment.

To compare side effects, t-tests were utilized. The analysis revealed no significant differences between the groups regarding the intensity of visual phenomena, itching, tingling, or burning sensations associated with tDCS (see Table 6). Only 3 participants reported experiencing at least one tDCS-related side effect post-wakening, likely due to sensations from the EEG cap and the conductive gel applied, as the participants were asleep during tDCS. The following day, no significant differences were observed between the groups in reports of skin redness, headache, fatigue, concentration difficulties, or sleep disturbances.

| | ***tDCS (M±SD)*** | ***Sham (M±SD)*** | ***Analysis*** |
|---|---|---|---|
| ***During tDCS*** | | | |
| **Visual phenomena** | 0 | .19 ± .75 | t(15) = -1, p = .333 |
| **Itching** | .125 ± .5 | .063 ± .25 | t(30) = .447, p = .658 |
| **Tingling** | .063 ± .25 | .125 ± .5 | t(30) = -.447, p = .658 |
| **Burning** | .063 ± .25 | 0 | t(15) = 1, p = .333 |
| **Pain** | 0 | 0 | / |
| ***After tDCS*** | | | |
| **Skin redness** | .125 ±.5 | 0 | t(15) = 1, p = .333 |
| **Headache** | .063 ±.25 | 0 | t(15) = 1, p = .333 |
| **Fatigue** | .5 ± 1.095 | .313 ± .873 | t(30) = .535, p = .596 |
| **Difficulties in concentration** | .313±.873 | .063 ± .25 | t(30) =1.101, p = .286 |
| **Nervousness** | 0 | 0 | / |
| **Sleeping disturbances** | .125 ± .5 | 0 | t(15) = 1, p = .333 |

Table 9. Post-stimulation side effect. Values presented are means (M) and standard deviations (SD). tDCS = transcranial direct current stimulation.
